# Advancing antimicrobial resistance monitoring in surface waters with metagenomic and quasimetagenomic methods

**DOI:** 10.1101/2022.04.22.489054

**Authors:** Andrea Ottesen, Brandon Kocurek, Padmini Ramachandran, Elizabeth Reed, Seth Commichaux, Gunnar Engelbach, Mark Mammel, Sanchez Saint Fleurant, Shaohua Zhao, Claudine Kabera, Amy Merrill, Nathalie Bonin, Hannah Worley, Noelle Noyes, Christina Boucher, Patrick McDermott, Errol Strain

**Affiliations:** Center for Veterinary Medicine, Food and Drug Administration, Laurel, Maryland, United States of America; Center for Food Safety and Applied Nutrition, Food and Drug Administration, College Park, Maryland, United States of America; Department of Computer Information Science and Engineering, University of Florida, Gainesville, Florida, United States of America; Department of Veterinary Population Medicine, University of Minnesota, Falcon Heights, Minnesota, United States of America

## Abstract

Surface waters present a unique challenge for the monitoring of critically important antimicrobial resistance. Metagenomic approaches provide unbiased descriptions of taxonomy and antimicrobial resistance genes in many environments, but for surface water, culture independent data is insufficient to describe critically important resistance. To address this challenge and expand resistome reporting capacity of antimicrobial resistance in surface waters, we apply metagenomic and quasimetagenomic (enriched microbiome) data to examine and contrast water from two sites, a creek near a hospital, and a reservoir used for recreation and municipal water. Approximately 30% of the National Antimicrobial Resistance Monitoring System’s critically important resistance gene targets were identified in enriched data contrasted to only 1% in culture independent data. Four different analytical approaches consistently reported substantially more antimicrobial resistance genes in quasimetagenomic data compared to culture independent data across most classes of antimicrobial resistance. Statistically significant differential fold changes (p<0.05) of resistance determinants were used to infer microbiological differences in the waters. Important pathogens associated with critical antimicrobial resistance were described for each water source. While the single time-point for only two sites represents a small pilot project, the successful reporting of critically important resistance determinants is proof of concept that the quasimetagenomic approach is robust and can be expanded to multiple sites and timepoints for national and global monitoring and surveillance of antimicrobial resistance in surface waters.

## Introduction

From a One Health perspective, surface waters function as key environmental integrators. They receive human, agricultural, and wildlife input and provide that same water for human, agricultural, and wildlife sustenance. Industrial and agricultural chemicals, metals, food additives, antibiotics, and even non-antibiotic drugs, have all been shown to play influential roles in the spread of AMR [1, 2]. Understanding the presence of pathogens and antimicrobial resistance (AMR) determinants in surface waters helps to inform risks across a wide range of applications and provides an integrative approach to public health. Recognizing the significant health impact that environmental water has on humans, animals, and the environment[3–5] the National Antimicrobial Resistance Monitoring System (NARMS) is investigating surface waters as a potential environmental modality for One Health AMR monitoring. This strategy requires methodological approaches capable of reporting AMR from an ecosystem as complex and dilute as water. Currently, NARMS monitoring efforts rely on standard in vitro antimicrobial susceptibility testing (AST) methods to generate minimum inhibitory concentrations (MICs) and the use of whole genome sequencing (WGS) to produce genotypes of single isolates. WGS data is currently being used to predict resistant phenotypes (without ASTs) and machine learning approaches are advancing MIC predictions directly from nucleotide data [6, 7]. These approaches rely on preliminary selective enrichment methods and produce excellent highly focused data characterizing single phenotypes of pathovars. Metagenomic data is useful for providing ‘big pictures’ of environmental, human, and animal microbiomes; however, for the species under active surveillance by NARMS, metagenomic data usually cannot provide sufficient depth of coverage to describe AMR phenotypes from complex matrices. Quasimetagenomic data (QMGS), due to its inclusion of enrichment, provides coverage of critically important resistance determinants with a broader throughput than culture independent (CI) metagenomics and WGS. Critically important antimicrobials (CIA) are ranked by the World Health Organization (WHO) according to their importance in human medicine in efforts to develop risk management for control of AMR in humans and animals. CIAs from WHO comprise more than a dozen different classes of antibiotics (Table 1) [8]. Currently NARMS monitors a subset of WHO CIAs derived from *Escherichia*, *Salmonella, Campylobacter* and *Enterococcus* primarily isolated from human, food-producing animals, raw retail meats, and feed environments. NARMS seafood monitoring also tracks resistance in *Aeromonas* and *Vibrio* species. In NARMS, critically important gene targets for monitoring include those conferring resistance to aminoglycosides, quinolones, ß-lactams, colistin, macrolides and ketolides, oxazolidinones, penicillins, etc. (Table 2) [9].

**Table 1.** World Health Organization Critically Important Antimicrobials (WHO CIAs)

|  |  |
| --- | --- |
| Aminoglycosides | Gentamicin |
| Ansamycins | Rifampin |
| Carbapenems and other penems | Meropenem |
| Cephalosporins (third and fourth generation) | Ceftriaxone |
| Phosphonic acid derivatives | Fosfomycin |
| Glycopeptides | Vancomycin |
| Glycylcyclines | Tigecycline |
| Lipopeptides | Daptomycin |
| Macrolides and ketolides | Erythromycin, telithromycin |
| Monobactams | Aztreonam |
| Oxazolidinones | Linezolid |
| Penicillins (natural, aminopenicillins, and antipseudomonal) | Ampicillin |
| Polymyxins | Colistin |
| Quinolones | Ciprofloxacin |
| Drugs used solely to treat tuberculosis or other mycobacterial diseases | Isoniazid |

**Table 2.** Genes Encoding Resistance to Critically Important Antimicrobial Agent.

| Macrolides | B-lactams | Colistins | Fluoroquinolones |
| --- | --- | --- | --- |
| 23S A2075G | *bla*<sub>CMY-2, 4, 30, 43,</sub> | *mcr-1.1, 1.2,*<br>3.1, 3.24 | *aac(6')-Ib-cr* |
| *acrB* R717L | *bla*<sub>CTX-M1, 2, 3, 8, 9, 14, 15, 27,</sub><br>32, 55, 65, 101, 104, 165, 190 |  | *qnrA1, B1, B2, B4,*<br>B6, B7, B9, B19,<br>B38, S1, S2, S4,<br>S13, VC1 |
| *ere(A)* | *bla*<sub>FOX-5</sub> |  | *gyrA* * |
| *erm 42, A, B, T* | *bla*<sub>KPC-2, 4</sub> |  | *gyrB* S464F |
| *mefB, C* | *bla*<sub>NDM-1</sub> |  | *msr(C)* |
| *mph A, B, E, G* | *bla*<sub>SHV-7,12, 30</sub> |  | *oqxA, B* |
| *msr(E)* | *bla*<sub>TEM-15, 207</sub> |  | *parC* ** |
|  |  |  | *parE* *** |
|  |  |  | *qepA, A1* |

Genes from *Salmonella*, *Escherichia coli*, *Campylobacter*, and *Enterococcus* are reported together.
\*D87G, D87N, D87Y, D90Y, P104S, S83A, S83F, S83L, S83Y, T86A, T86I, T86K, T86V
\*\**parC* A56T, E84G, S801I, S80R,
\*\*\**parE* D475E, I529L, L416A

For NARMS’s expanded One Health mission to effectively examine AMR in surface waters, new approaches will be required to describe critically significant resistomes. In the present study dead end ultrafiltration[10] was the water collection method used to collect 10L and 50L sample volumes from each of two sites: 1) an urban creek in close proximity to a hospital (Sligo), and 2) a recreational reservoir which is the source of drinking water for Prince George’s county, Maryland (Patuxent). DNA extracted from water was evaluated using both metagenomic and quasimetagenomic data to assess the ability of each data type to describe critically important NARMS resistance determinants. Quasimetagenomics, which uses shotgun sequencing of enriched microbiomes at strategic temporal increments during pathogen recovery has previously proven useful for expedited source tracking in outbreak investigations [11], identification of multi-serovar diversity associated with samples linked to outbreaks, and identification of co-competitors, co-enrichers, and recovery biases in state of the art FDA microbiological culturing methods [12, 13]. Here we demonstrate how the approach also greatly enhances the capacity for resistome reporting and provides data which can differentiate AMR by water source, detect close to 30% of NARMS critically important resistance determinants, detect taxa important to global AMR mortality, and potentially predict and identify emerging resistance.

## Results

### Resistome Composition of the Sligo and Patuxent Water by CI and QMGS approaches

Results from all four bioinformatic pipelines and corresponding databases (AMRFinder Plus[14], COSMOSID [15], AMR++ 2.0 [16], and CARD [17] demonstrated that data from QMGS data greatly expanded the capacity for resistome reporting for Sligo and Patuxent water. Fig 1 shows annotations of ß-lactams by AMR++ with normalization[16], AMRFinder Plus, CARD[17] and COSMOSID (www.cosmosid.com) for both CI and QMGS samples. No ß-lactam annotations (using default parameters and normalization) were reported by any of the pipelines for CI water samples except by the kmer based COSMOSID pipeline. (Kmer based approaches do have well demonstrated sensitivity, but without dedicated curation and validation, specificity can be unreliable). Reporting for almost every class of drug, biocide, metal, and multi-compound resistance was vastly increased using the QMGS approach (Figs 1 and 2 and S1 Fig). With CI data alone, it was not possible to evaluate differences between the two sites (Sligo and Patuxent) for trimethoprims, sulfonamides, phenicols, oxazolidinones, nucleosides, metronidazoles, lipopeptides, glycopeptides, fosfomycins, ß-lactams, bacitracins, and many other groups due to lack of coverage in at least one set of CI samples and often both (Fig 3).

**Fig 1.**
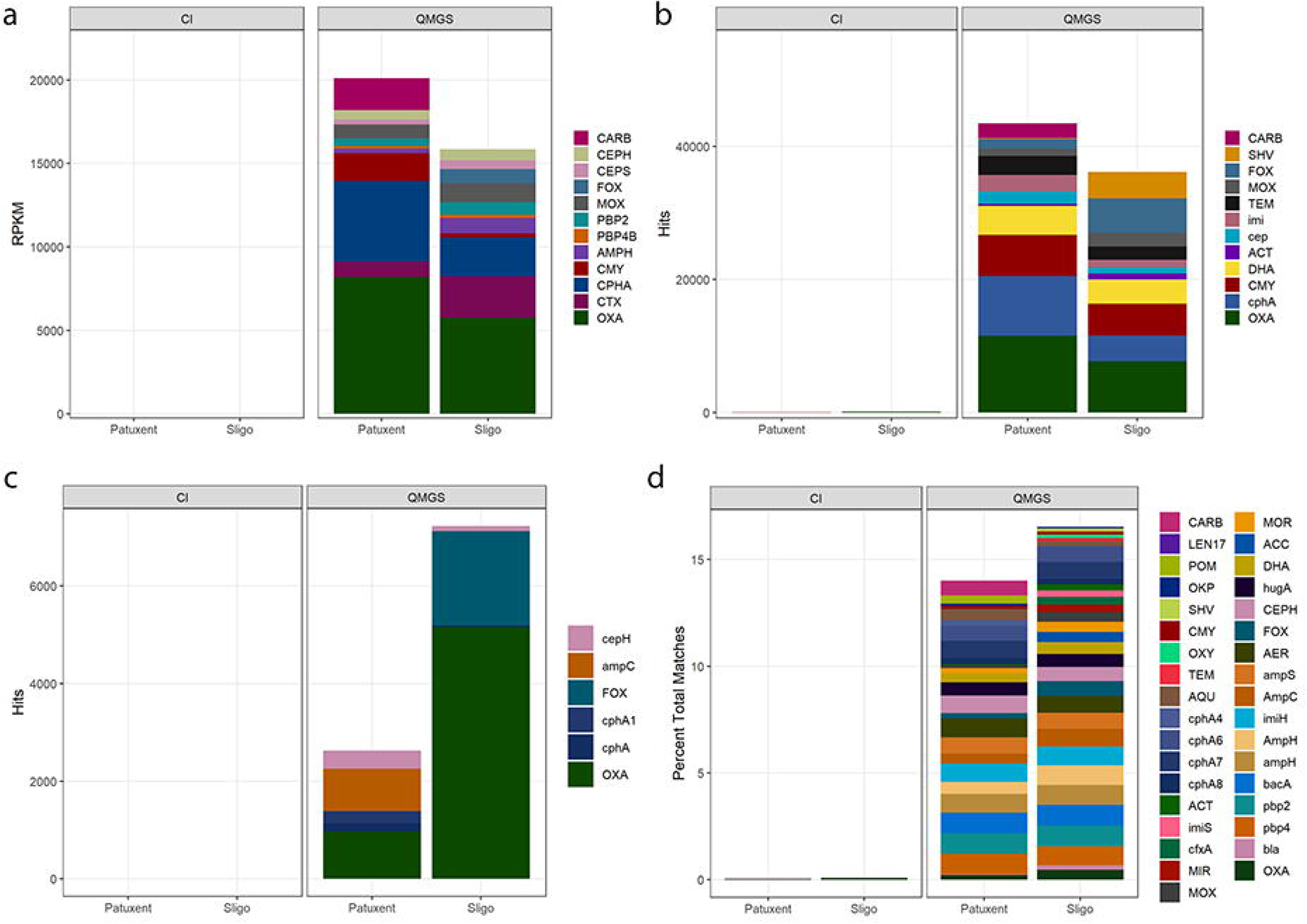
Resistome Composition: ß-lactams by different annotation pipelines. Four different analytical approaches, a) AMR++, b) CARD, c) AMRFinder, and d) COSMOS ID, consistently reported substantially more ß-lactam antimicrobial resistance genes in QMGS data compared to CI data across almost all classes of antimicrobial resistance.

**Fig 2.**
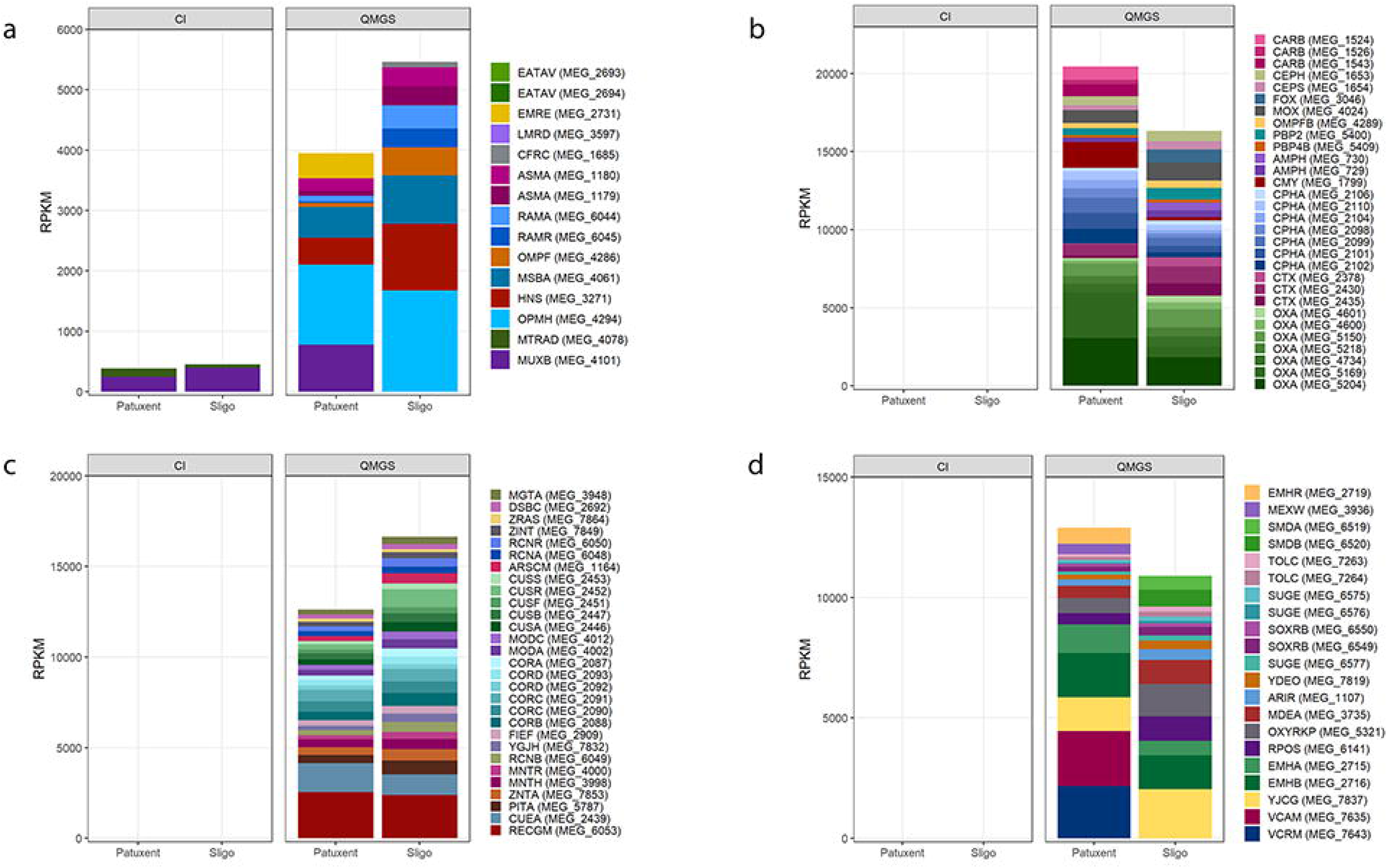
Resistome Composition: Sligo and Patuxent Water by CI and QMGS approaches. Detection of AMR genes in CI and QMGS data are shown for a) multidrug resistance b) ß-lactams c) multimetal resistance and d) multibiocide resistance. Multidrug resistant genes: MTRAD* (some of which require SNP validation) and *muxB* were the only genes detected in culture independent data. All other gene coverage for multidrug resistance, ß-lactams, multimetal resistance, and multibiocide resistance was only observed in QMGS data.

**Fig 3.**
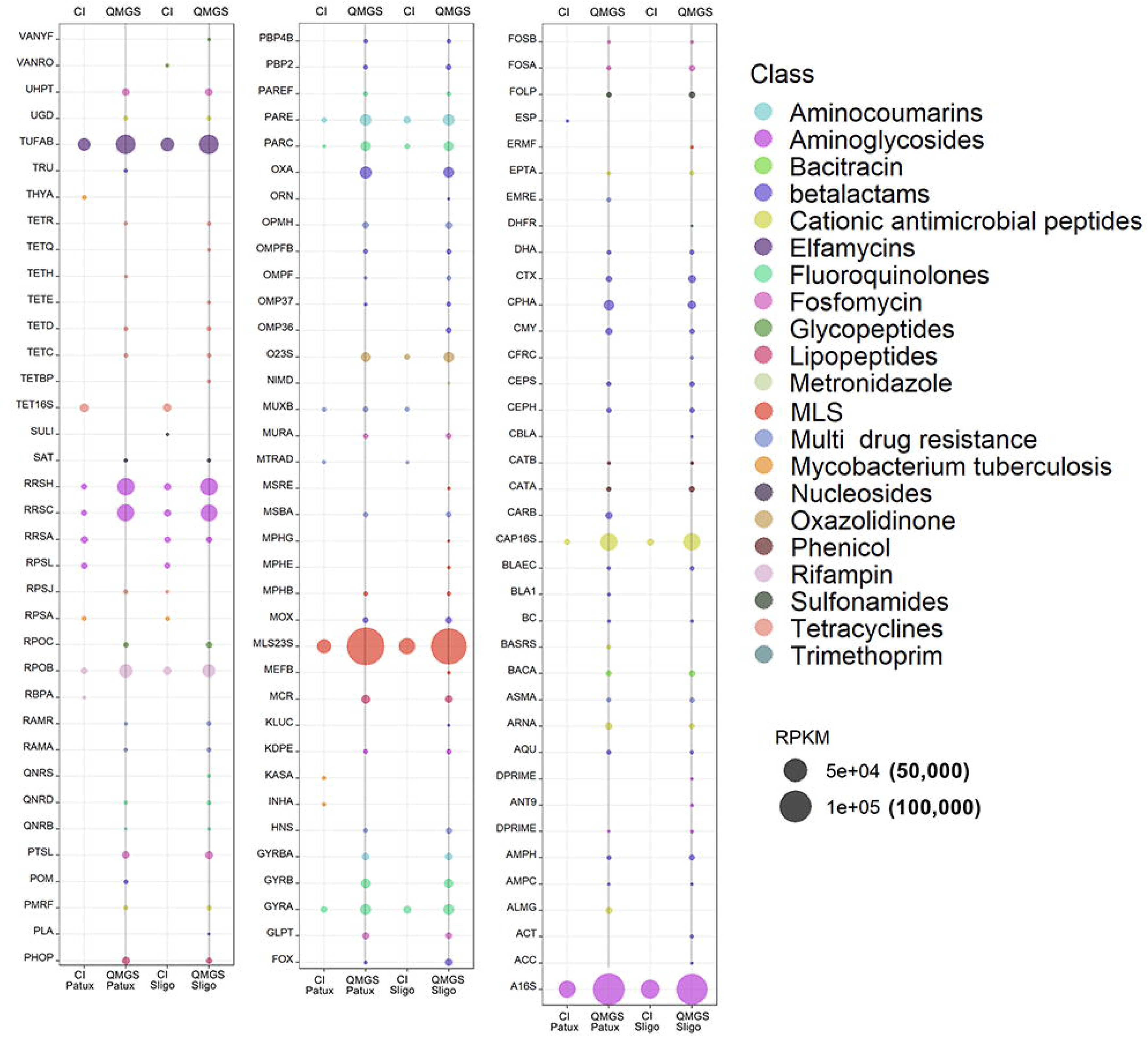
Resistome Composition: Drugs by Gene in Sligo and Patuxent (CI&QMGS). Overview of antimicrobial resistance to drugs observed in Patuxent and Sligo water for culture independent (CI) and quasimetagenomic (QMGS) samples by gene. The differences between CI and QMGS are shown for genes of major classes of AMR. (A darker line runs through the QMGS samples to make it easy to differentiate QMGS data from CI data). Many more gene hits were observed in QMGS samples. With the exception of a very few genes some of which require SNP validation, TET16S, *Mycobacterium tuberculosis* specific *rpsA* and MTRAD*, the data from QMGS provided significantly more expansive coverage of ARGs.

### Resistome Composition: Identification of NARMS critically important antimicrobial resistance

A key objective for this work was to evaluate metagenomic and quasimetagenomic data for their potential contribution to detection of resistance genes important to NARMS monitoring. From the list of NARMS critically important resistance gene targets, more than 30% were identified across the two water sources using the QMGS approach compared to only 1% identified using metagenomics. Critically important gene targets were observed at 100% identity across 100% coverage using the Comprehensive Antibiotic Resistance Database (CARD) for Sligo and Patuxent QMGS data; *bla*_TEM-207_, *bla*_TEM-15_, *bla*_SHV-30_, *bla*_SHV-12_, *qnrS2, qnrB9, qnrB7, qnrB6, qnrB2, qnrB19, qnrB1, oqxB, oqxA, msrE, mphE, mphA, mefC*, *bla*_FOX-5_, *ereA*, *bla*_CMY-43_, *bla*_CMY-4_, *bla*_CMY-30_, *bla*_CMT-2_, *acrB*. With the exception of *ereA,* observed in CI Sligo water samples, all genes of importance to NARMS monitoring were observed only in QMGS samples (Fig 4).

**Fig 4.**
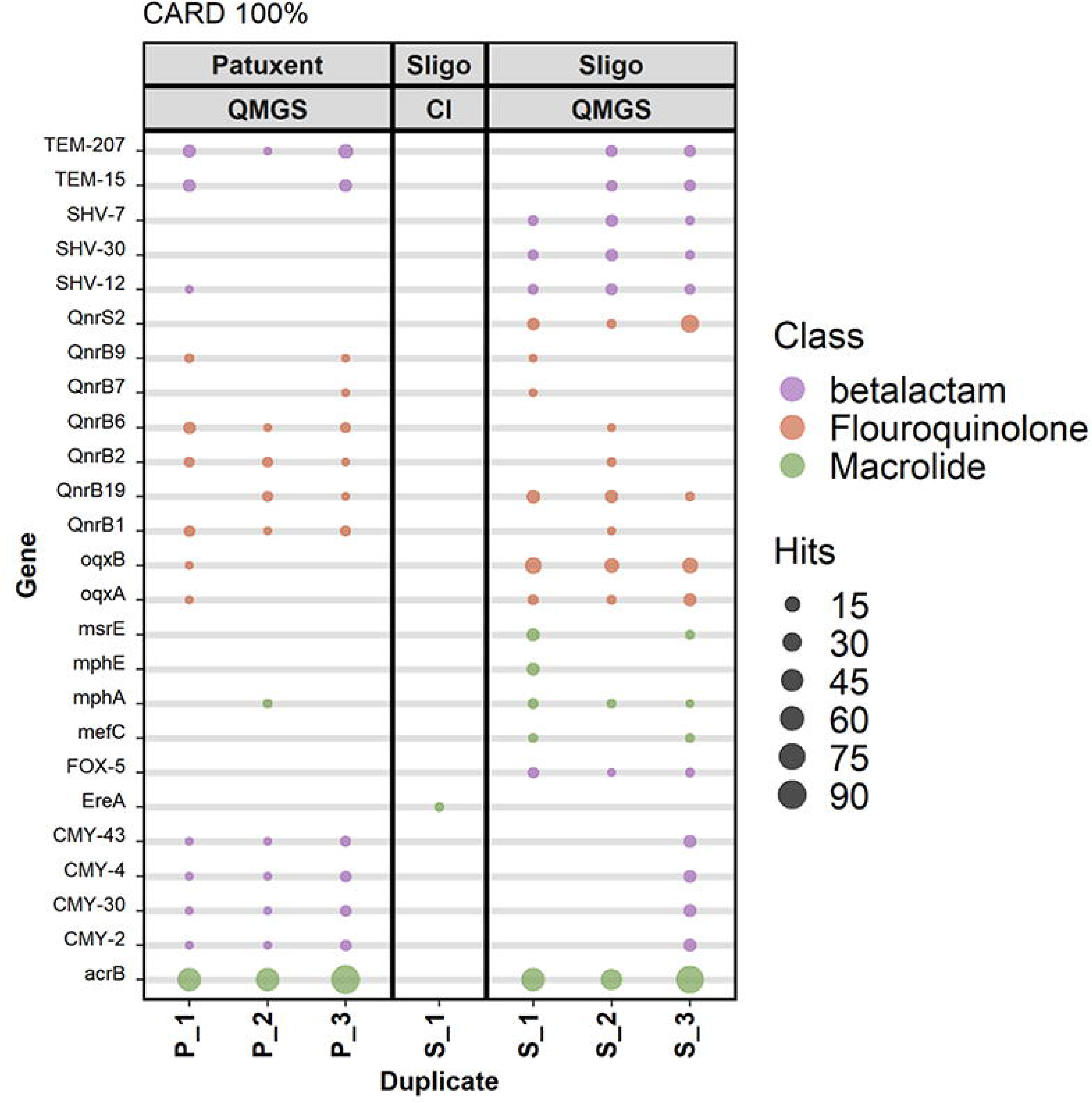
NARMS critically important genes. From the list of NARMS critically important antimicrobial (CIA) resistance genes, using 100% identity across 100% of gene length (CARD), almost 30% (29.4) were identified across the two water sources using the QMGS approach compared to 1% identified in CI samples.

### Resistome Composition: Quinolone Resistance Genes in Sligo and Patuxent Water

Fluoroquinolone resistance genes and variants were reported by the AMR++ pipeline (Megares 2.0 database) often with 100 percent identity across 100% of the gene length corresponding to published reports of resistant alleles. However, all observed MEGARes fluoroquinolone accessions included the addendum “*RequiresSNPConfirmation*”. Additional confirmation analytics were conducted ‘by hand’ comparing resistance determining alleles to genes identified in Sligo and Patuxent water to ensure validated annotation of fluoroquinolone resistance determinants [18]. From Patuxent water, the resistance determining allele of *gyrB* S464Y (1) was confirmed while all others were identified as wildtype alleles. In Sligo water there were quite a few more fluoroquinolone resistance determining alleles confirmed; including *gyrB* E466D (1), S464Y (1), *gyrA* S83N(2), S83T, and *parC* S80R, S80I. The AMRFinderPlus pipeline reported *qnrD, qnrD1*, and *qnrS2* but no other genes for quinolone or fluoroquinolones were identified in Sligo or Patuxent water samples using that pipeline.

### Resistome Composition: ß-lactams

As already indicated, no ß-lactams were detected in CI samples except for the few detected by the kmer based COSMOSID pipeline. The number of hits to ß-lactams in enriched QMGS samples however provided sufficient data to describe differences by site. While it is possible that some differences were artifacts of enrichment biases and not reflective of resistome biology, it is equally likely that genes with statistically significant fold changes (p < 0.05) may be reflective of true resistome abundances in Sligo and Patuxent waters. Fig 5 shows statistically significant fold changes (p< 0.05) in ß-lactams associated with Sligo and Patuxent water.

**Fig 5.**
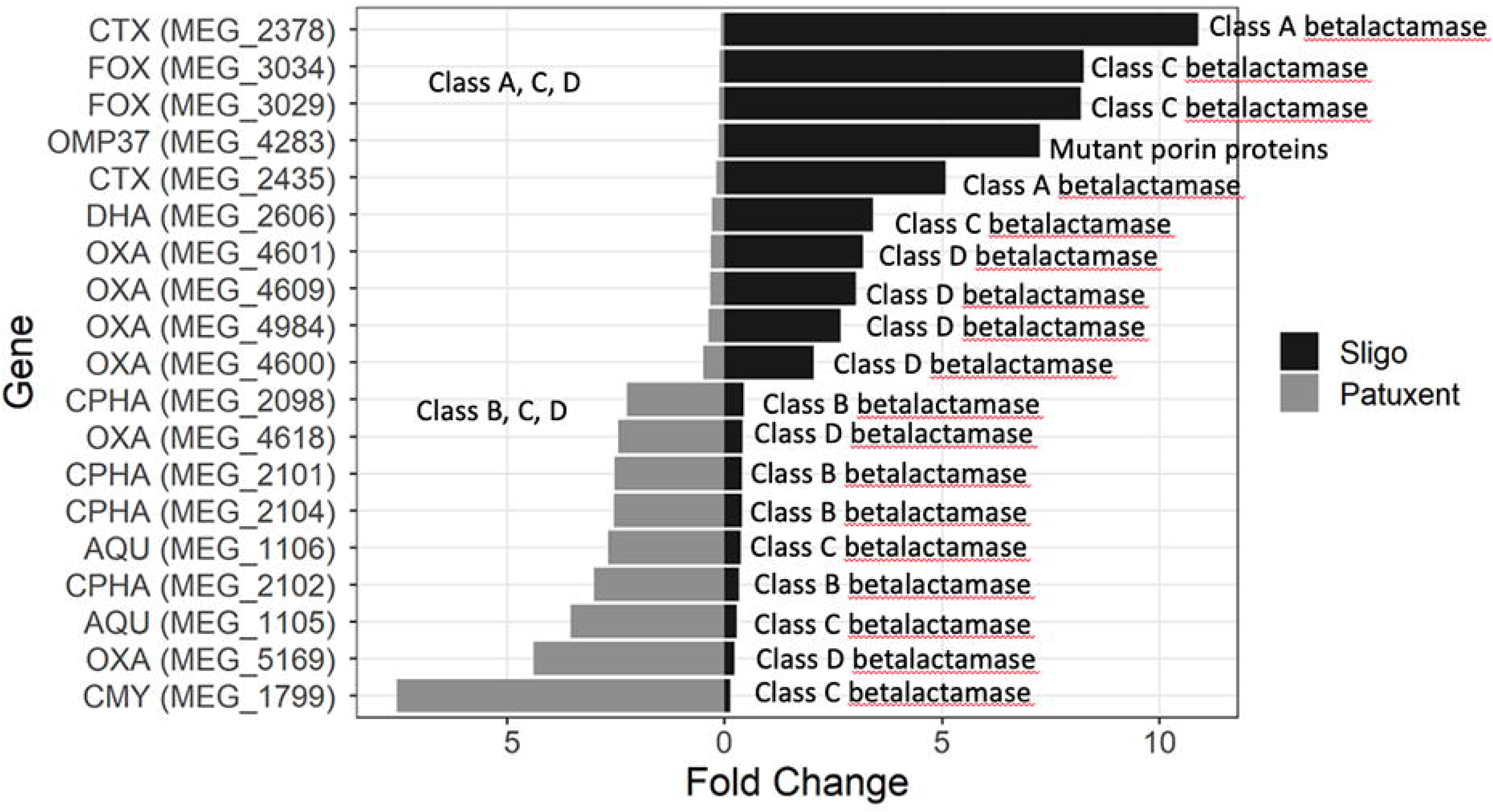
Differential Abundance of ß-lactams in Sligo and Patuxent Water by QMGS Data. Genes with statistically significant values (p< 0.05) for fold change differences from one water source to the other (Patuxent - Sligo) are shown.

### Resistome Composition: Plasmids

Though many plasmids were identified in CI data, none could be annotated by Platon, a pipeline for identification and characterization of bacterial plasmid contigs in short-read draft assemblies exploiting protein sequence-based replicon distribution scores [12]. In QMGS data however, for both Sligo and Patuxent, numerous plasmids were identified and annotated using Platon (Fig 6, Tables 3 and 4). IncFIIK, IncQ2, Col(pHAD28) were only observed in Sligo while others were observed in both Sligo and Patuxent. InC, Inc, and IncI plasmids often carry AMR genes especially in Enterobacteriaceae. IncHI and IncHII’s plasmid types are also frequently associated with AMR genes [19] and observed IncI plasmid (Gamma1AP005147) shares genomic similarity to the pESI megaplasmid found in *Salmonella* serovar Infantis.

**Fig 6.**
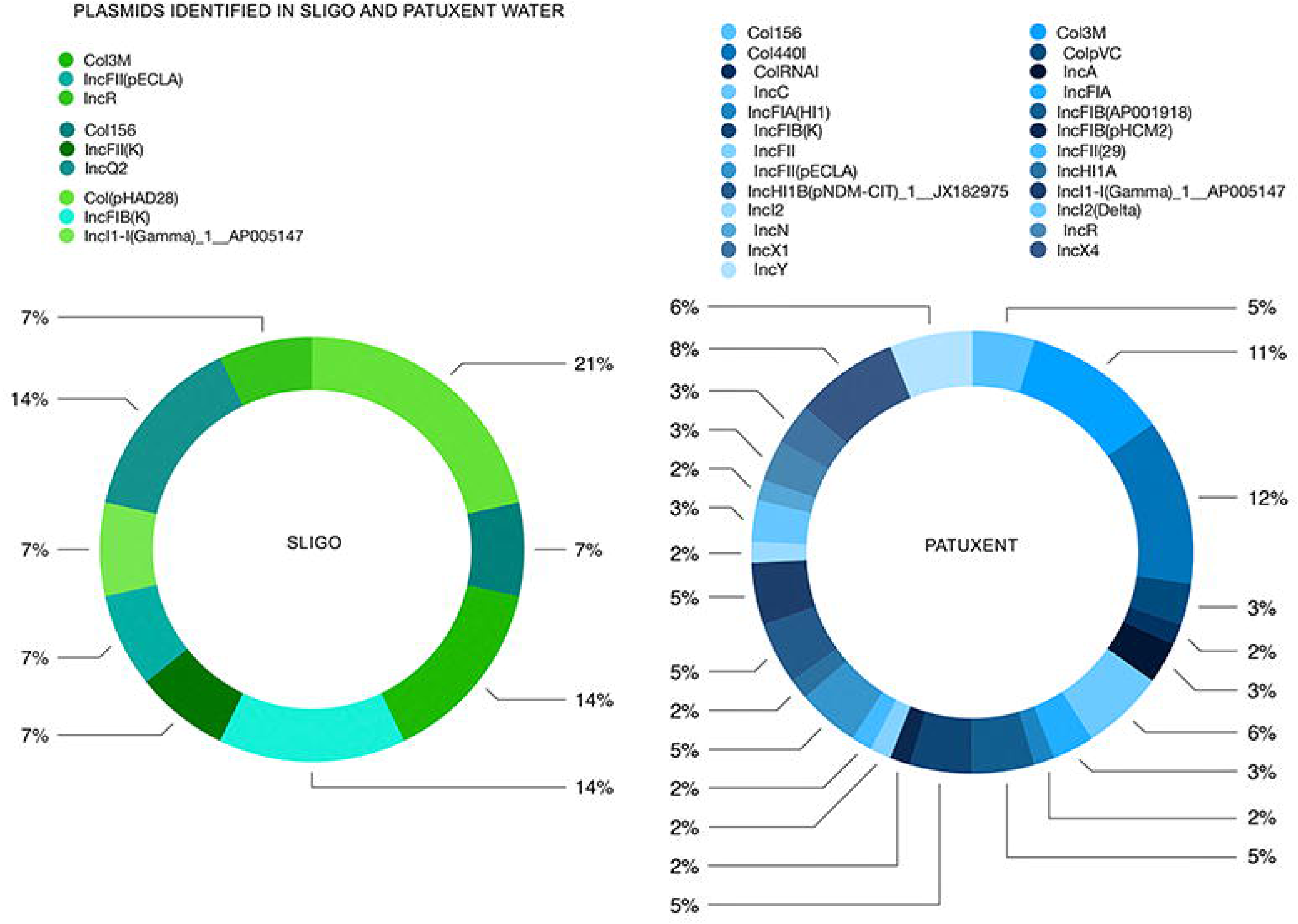
Plasmids in Sligo and Patuxent Water. Composition of plasmids that could be identified in QMGS data sets from Sligo and Patuxent water (none could be annotated from CI data although many were detected).

**Table 3.** Sligo Plasmids and Associated Taxonomy (Platon)

| Kraken_taxonomy | Plasmid_type2 |
| --- | --- |
| *Escherichia coli* ABU 83972 (taxid 655817) | Col(MG828) |
| *Shigella sonnei* (taxid 624) | Col(pHAD28) |
| Enterobacteriaceae (taxid 543) | Col(pHAD28) |
| *Escherichia coli* (taxid 562) | Col156 |
| *Proteus mirabilis* (taxid 584) | Col3M |
| *Proteus mirabilis* (taxid 584) | Col3M |
| *Klebsiella pneumoniae* (taxid 573) | IncFIB(K) |
| *Klebsiella variicola* (taxid 244366) | IncFIB(K) |
| *Raoultella planticola* (taxid 575) | IncFII(K) |
| *Enterobacter hormaechei* subsp. *steigerwaltii* (taxid 299766) | IncFII(pECLA) |
| *Escherichia coli* (taxid 562) | IncI1-I(Gamma)_1__AP005147 |
| *Chlamydia suis* (taxid 83559) | IncQ2 |
| unclassified (taxid 0) | IncQ2 |

**Table 4.** Patuxent Plasmids and Associated Taxonomy (Platon)

| Kraken_taxonomy | plasmid type |
| --- | --- |
| *Escherichia coli* ABU 83972 (taxid 655817) | Col(MG828) |
| *Escherichia coli* (taxid 562) | Col156 |
| *Proteus mirabilis* (taxid 584) | Col3M |
| Enterobacteriaceae (taxid 543) | Col440I |
| *Salmonella enterica* subsp. *enterica* serovar Typhimurium (taxid 90371) | Col440I |
| *Klebsiella pneumoniae* (taxid 573) | Col440I |
| *Escherichia coli* (taxid 562) | Col440I |
| *Enterobacter* sp. FY-07 (taxid 1692238) | Col440I |
| *Enterobacter hormaechei* subsp. *steigerwaltii* (taxid 299766) | Col440I |
| *Salmonella enterica* subsp. *enterica* serovar Enteritidis (taxid 149539) | ColpVC |
| Enterobacteriaceae (taxid 543) | ColRNAI |
| *Enterobacter hormaechei* subsp. *steigerwaltii* (taxid 299766) | IncA |
| *Enterobacter hormaechei* subsp. *steigerwaltii* (taxid 299766) | IncC |
| *Escherichia coli* O157:H7 (taxid 83334) | IncFIA |
| *Escherichia coli* (taxid 562) | IncFIA |
| *Klebsiella pneumoniae* (taxid 573) | IncFIA(HI1) |
| *Escherichia coli* (taxid 562) | IncFIB(AP001918) |
| *Escherichia coli* O104:H4 (taxid 1038927) | IncFIB(AP001918) |
| *Klebsiella variicola* (taxid 244366) | IncFIB(K) |
| *Klebsiella pneumoniae* (taxid 573) | IncFIB(K) |
| Enterobacteriaceae (taxid 543) | IncFIB(pHCM2) |
| *Escherichia coli* (taxid 562) | IncFII |
| *Escherichia coli* (taxid 562) | IncFII(29) |
| *Enterobacter* sp. N18-03635 (taxid 2500132) | IncFII(pECLA) |
| *Salmonella enterica* subsp. *enterica* serovar Typhi (taxid 90370) | IncHI1A |
| *Klebsiella oxytoca* (taxid 571) | IncHI1B(pNDM-CIT)_1__JX182975 |
| *Escherichia coli* (taxid 562) | IncI1-I(Gamma)_1__AP005147 |
| *Escherichia coli* W (taxid 566546) | IncI1-I(Gamma)_1__AP005147 |
| *Escherichia coli* (taxid 562) | IncI2 |
| *Escherichia coli* (taxid 562) | IncI2(Delta) |
| *Salmonella enterica* (taxid 28901) | IncN |
| *Escherichia coli* (taxid 562) | IncR |
| Enterobacteriaceae (taxid 543) | IncR |
| *Salmonella enterica* subsp. *enterica* serovar Heidelberg (taxid 611) | IncX1 |
| *Escherichia coli* (taxid 562) | IncX1 |
| *Escherichia albertii* (taxid 208962) | IncX4 |
| Enterobacteriaceae (taxid 543) | IncX4 |
| *Escherichia coli* (taxid 562) | IncX4 |

While the vast majority of antimicrobial resistance genes were identified in QMGS data, there was a small number of genes for which the converse was true; *i.e.,* more information was provided by culture independent (CI) assessment than by QMGS. Almost half of the genes observed exclusively in CI data (AMR++) were *Mycobacterium tuberculosis* drug specific, although each annotation had the addendum that further confirmation would be needed to confirm resistance-determining single nucleotide polymorphisms (SNPs). SNP validation is currently in development for *Mycobacterium tuberculosis* resistance determinants but the observation that *Mycobacterium* spp. were almost exclusively observed in culture independent data will be important for environmental *Mycobacterium* surveillance efforts. The length of time needed to culture a *Mycobacterium* isolate has been described as 6 days or more [20] so it is not surprising that a 24H timepoint in a broth designed primarily for enrichment of Enterobacteriales would not provide the right conditions to observe *Mycobacterium* populations.

### Microbiome Composition: Bacteria from Sligo and Patuxent by CI & QMGS approaches

Diverse taxonomic compositions were observed between the CI Sligo and Patuxent water microbiota; Burkholderiales, Comamonadaceae, Aeromonadales, and Caulobacterales were abundant in Sligo water but not in the Patuxent samples. Actinobacteria, Alphaproteobacterium, and Chroococcales were abundant in Patuxent water and not in Sligo (CI data shown in Fig 7 a & b). Sligo and Patuxent QMGS taxonomy was much more homogenous (QMGS data shown in Fig 7 c & d). Enterobacteriales were present in less than 1% of CI samples but comprised between 15% and 20% of QMGS water enrichments. After enrichment, samples are obviously biased toward recovery of taxa that thrive in nutrient conditions, in this case BPW, comprised of peptone, sodium chloride, disodium phosphate, mono-potassium phosphate and distilled water, and TSB, comprised of trypticase peptone, phytone peptone, sodium chloride, dipotassium phosphate, glucose and distilled water [21]. While enrichments obscure certain taxonomic and functional features of the true microbiome, they also provide robust data to surveil critically important AMR from medically relevant taxa.

**Fig 7.**
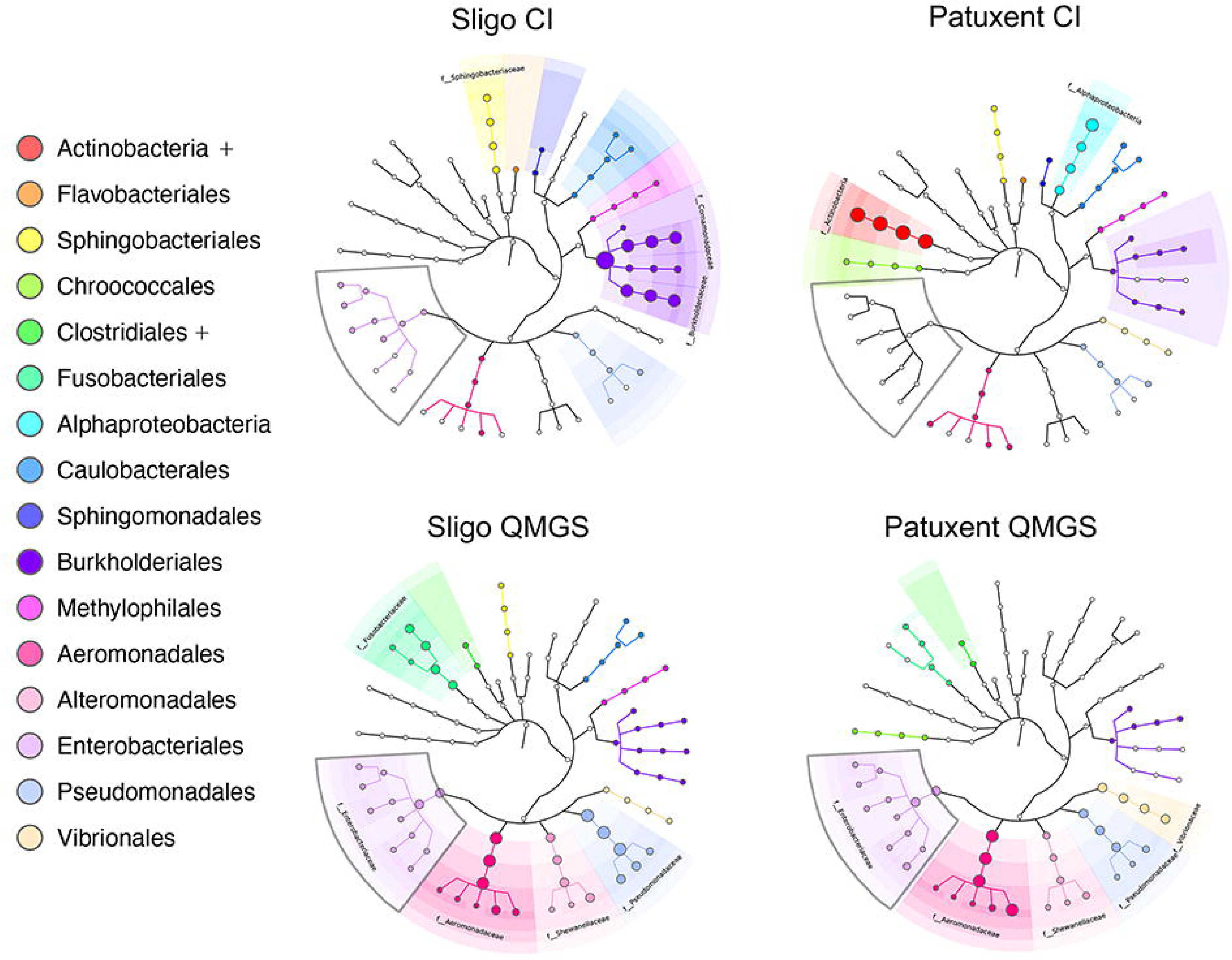
Microbiome Composition: Bacterial Taxa from Sligo and Patuxent CI and QMGS. Enterobacteriales were present in less than 1% of CI samples (a & b) but comprised between 15% and 20% of QMGS water enrichments (c & d). Shaded regions are highlighted if they correspond to taxa with >1% abundance in a sample. If the abundance is less than 1 percent, there is no area shading but a node may still have a color when observed at less than 1 percent.

### Critically Important Antimicrobial Resistance

The Sligo and Patuxent data were examined for key NARMS monitoring targets; *Campylobacter, Salmonella, Escherichia coli*, *Enterococcus,* and NARMS seafood targets; *Aeromonas* and *Vibrio* (Fig 8). Data were also examined for presence of ‘ESKAPE’ nosocomial pathogens, known for carriage of multidrug resistance and virulence. These are *Enterococcus faecium, Staphylococcus aureus, Klebsiella pneumoniae, Acinetobacter baumannii, Pseudomonas aeruginosa, and Enterobacter spp.,* and for pathogens with the highest number of global deaths attributable to AMR; *Escherichia coli*, *Staphylococcus aureus*, *Klebsiella pneumoniae*, *Streptococcus pneumoniae*, *Acinetobacter baumannii*, and *Pseudomonas aeruginosa* [22]. ESKAPE bacteria and bacteria attributable to the highest numbers of AMR associated deaths, were combined to create a list of 8 important bacterial monitoring targets: *Staphylococcus aureus, Klebsiella pneumonia, Acinetobacter baumannii, Pseudomonas aeruginosa*, *Escherichia coli, Streptococcus pneumoniae, Enterococcus faecium* and *Enterobacter* spp. (Fig 9). Figs 8 and 9 show hits to target species in the first panel and hits to all observed species from same genera in the second panel. This comprehensive ecological view of all species in the genus of a surveillance target may prove useful for identification of emerging pathogens, resistance, and risk assessment.

**Fig 8.**
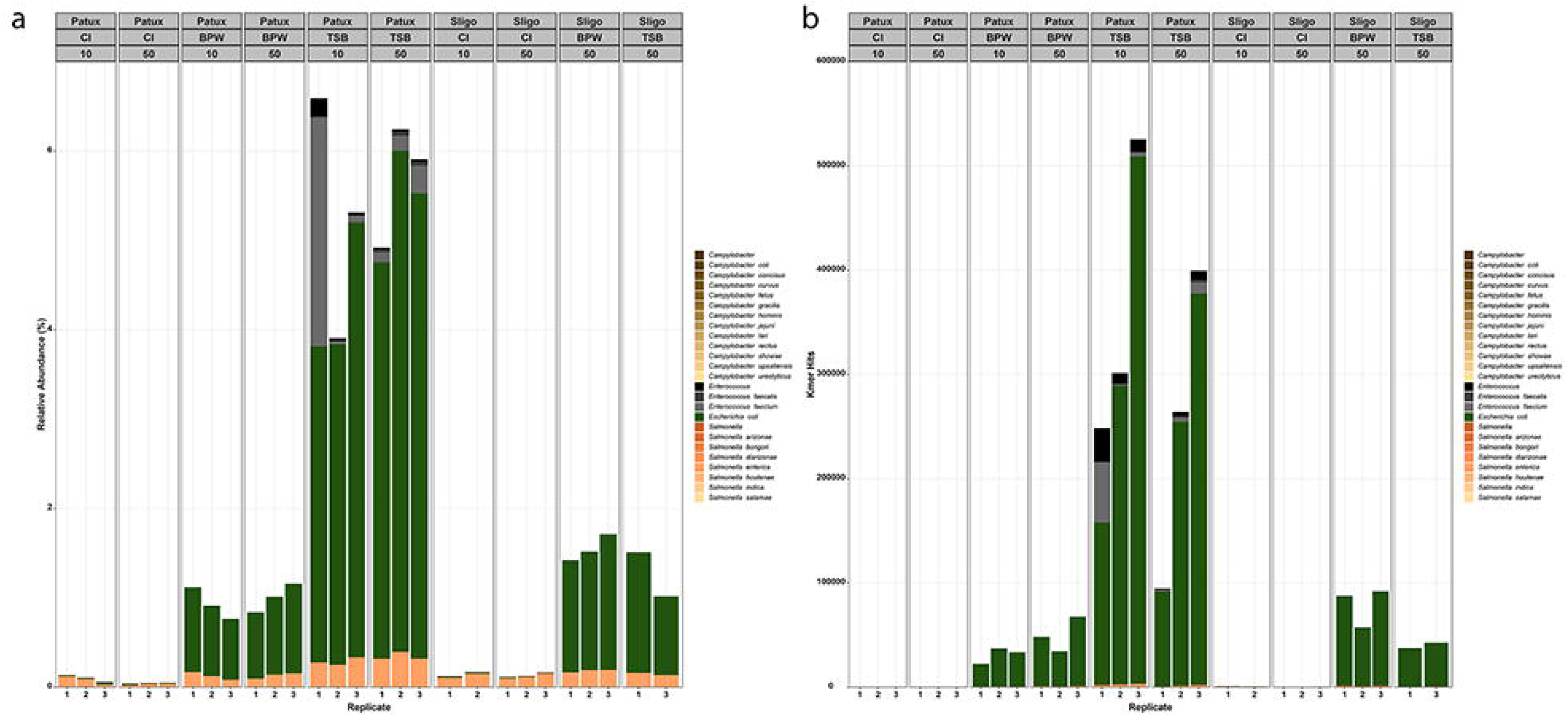
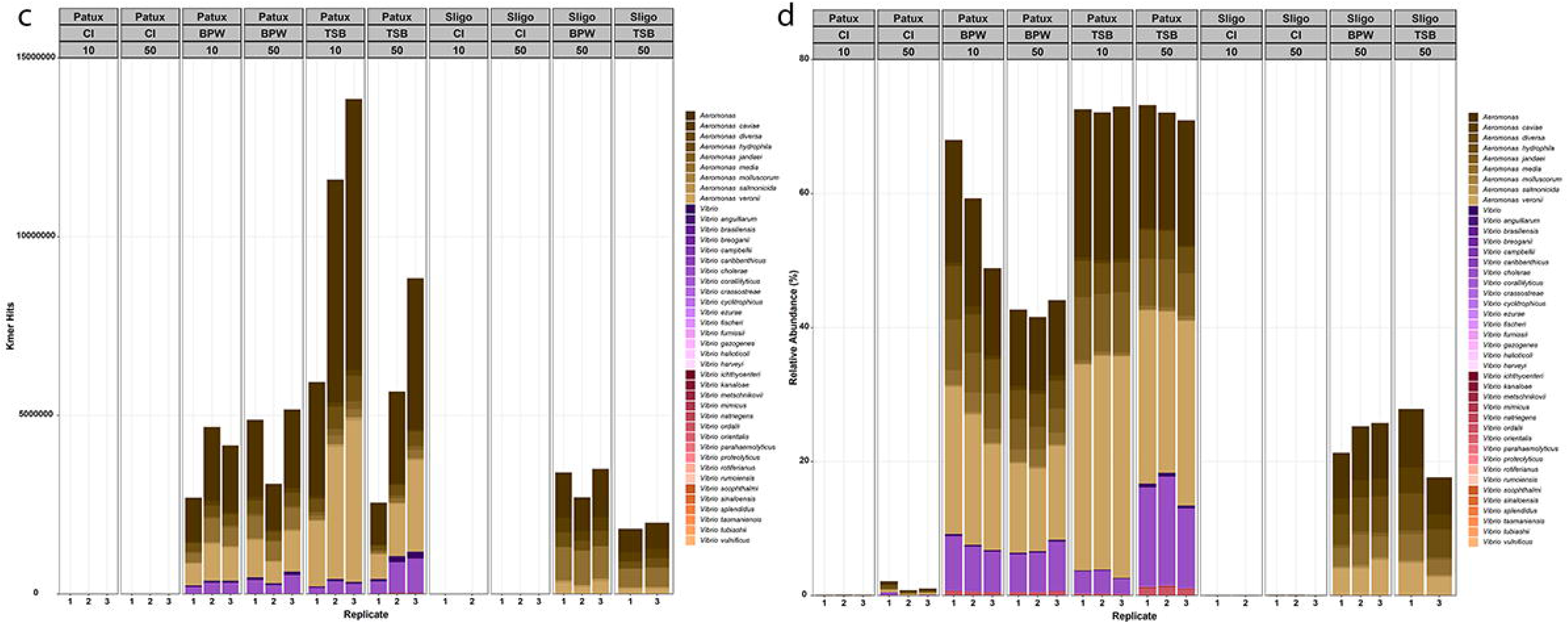
Species important to NARMS Monitoring. Panel A shows NARMS targets; *Campylobacter, Salmonella, Enterobacter and Escherichia* genera and all related species by relative abundance contrasted to Panel B which shows kmer hits. Panel C and D show relative abundance and kmer kits for NARMS seafood pathogens *Vibrio* and *Aeromonas* and associated species.

**Fig 9.**
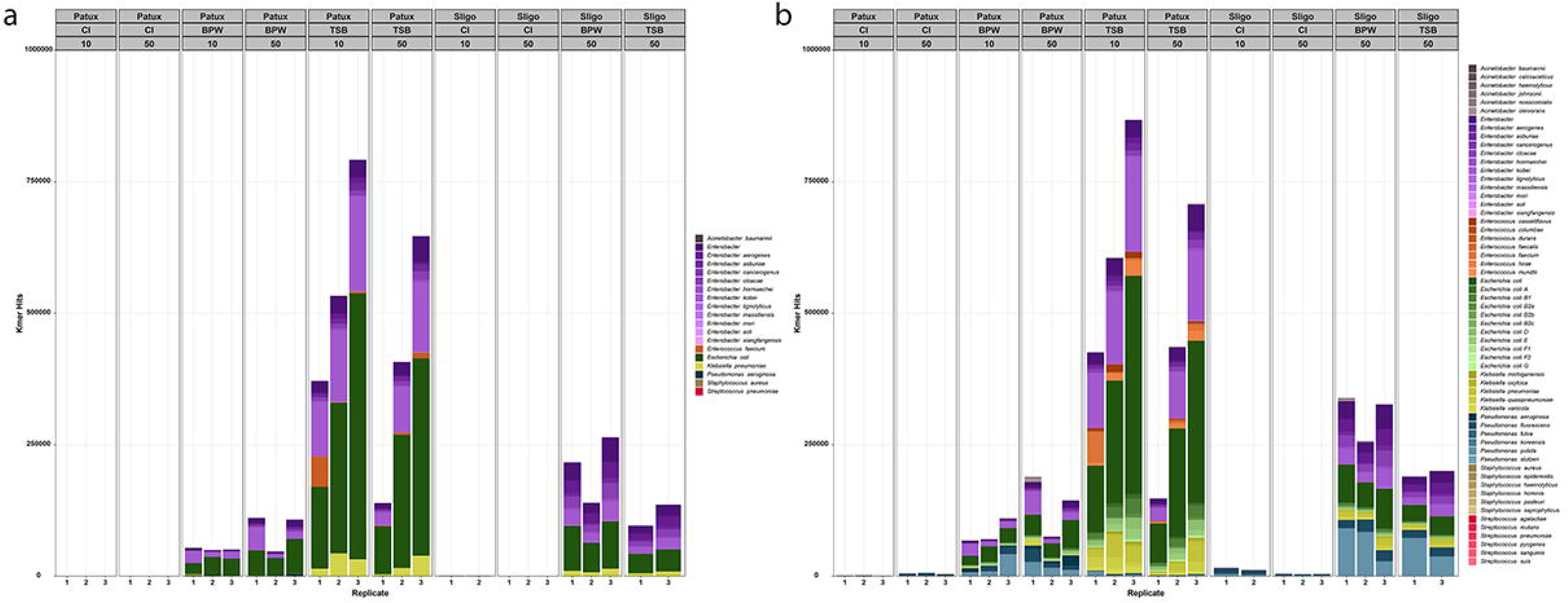
ESKAPE8 pathogens. The ‘ESKAPE8’ list used here is comprised of the 6 ESKAPE pathogens with the only two not represented from the list of top 6 species responsible for global AMR associated mortality. In panel A the genus and species of pathogens under active surveillance is shown and in panel b) all the close relatives – other species within the genera are shown.

### Bacterial Taxa of Importance to Veterinary Monitoring Efforts

Important veterinary pathogens were also detected from Sligo and Patuxent water samples. The Veterinary Laboratory Investigation and Response Network (Vet-LIRN) regularly monitors *E. coli* and *Salmonella* (NARMS targets, Figure 8) and the *Staphylococcus intermedius* group (SIG). The SIG, comprised of *Staphylococcus pseudointermidius, Staphylococcus intermedius, and Staphylococcus delphini,* is a collective of emerging importance [23]. The European Antimicrobial Resistance Surveillance network in Veterinary medicine (EARS-Vet) surveils 11 bacterial targets; *Escherichia coli, Klebsiella pneumoniae, Mannheimia haemolytica, Pasteurella multocida, Actinobacillus pleuropneumoniae, Staphylococcus aureus, Staphylococcus pseudintermedius, Staphylococcus hyicus, Streptococcus uberis, Streptococcus dysgalactias* and *Streptococcus suis,* across six animal species (cattle, swine, chickens (broilers and laying hens) turkeys, cats, and dogs)[24]. Genera and species from EARS-Vet surveillance that were observed in Sligo and Patuxent water are shown in Fig 10. As previously indicated, most pathogen species under active surveillance require general and selective enrichment protocols before they can be detected, sequenced and described. Certain genera, however, are more robustly reported by culture independent data. *Mycobacterium, Neisseria, Mycoplasma*, and *Borrelia* for example, were predominantly observed in CI data from Sligo and Patuxent water compared to QMGS data (Fig 11).

**Fig. 10.**
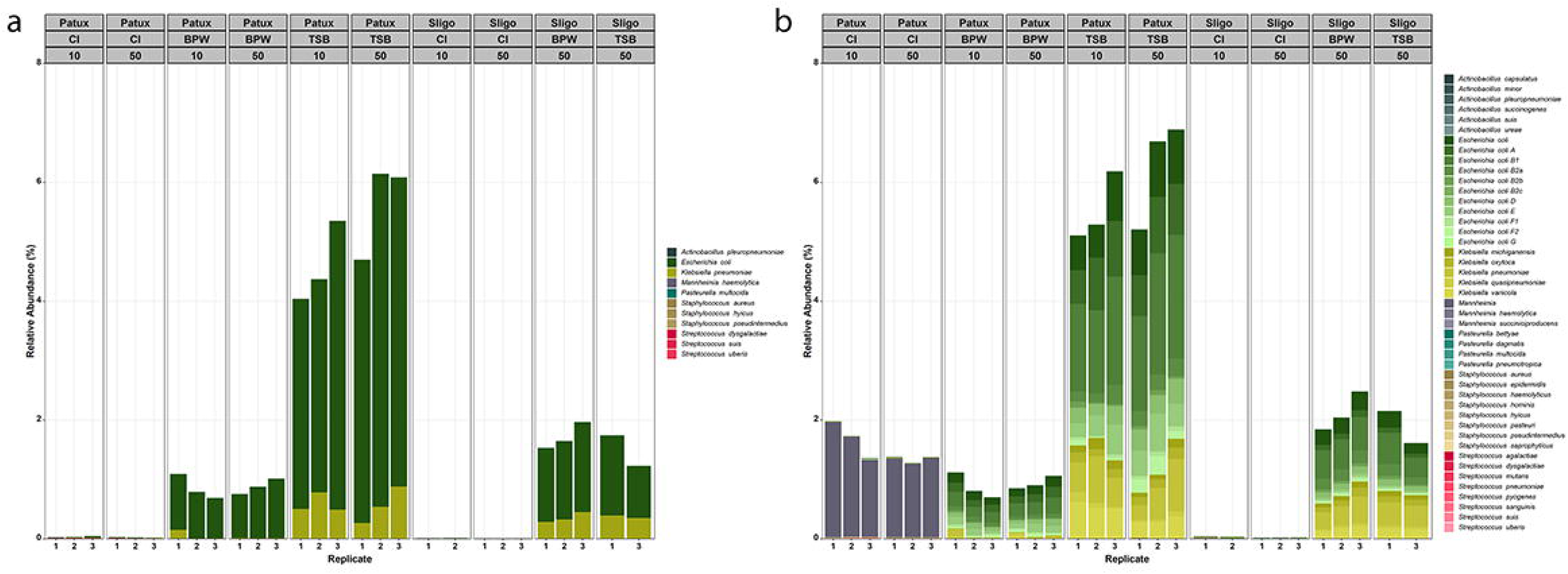
**EARS-Veterinary Surveillance Targets**. The European Antimicrobial Resistance Surveillance network in Veterinary medicine (EARS-Vet) surveils 11 bacterial targets; *Escherichia coli, Klebsiella pneumoniae, Mannheimia haemolytica, Pasteurella multocida, Actinobacillus pleuropneumoniae, Staphylococcus aureus, Staphylococcus pseudintermedius, Staphylococcus hyicus, Streptococcus uberis, Streptococcus dysgalactias* and *Streptococcus suis,* across six animal species (cattle swine, chickens (broilers and laying hens) turkeys, cats and dogs). Incidence of these taxa is shown for Sligo and Patuxent water.

**Fig 11.**
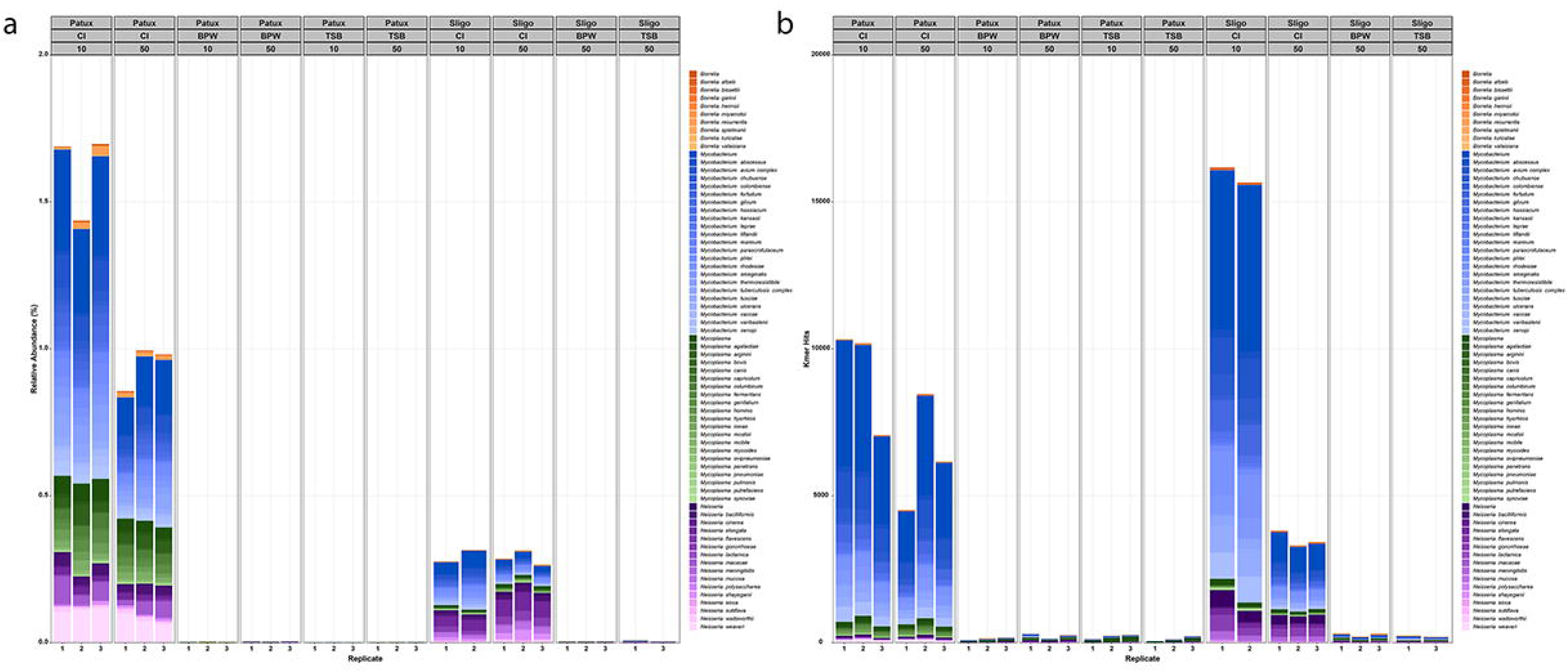
Incidence of *Borrelia, Mycobacterium, Mycoplasma* and *Neisseria* in Sligo and Patuxent water is primarily associated with CI data. As previously described for *Mycobacterium*, it is important for monitoring efforts to understand when CI data will be more useful than QMGS data and vice versa. All the species shown here were primarily observed in CI data.

## Discussion

Water plays perhaps the most important role in states of health and disease in humans and other animals, yet it is one of the most complex matrices to monitor due to its dilute nature. Work by Gweon et al. used 200 million CI reads to describe AMR in pig caeca, effluent, and stream sediment. For pig caeca, hits to AMR genes reached about 50,000 and for effluent, about 22,000; for stream sediment only 22 hits were observed for the most abundant AMR genes[25]. This is consistent with numbers observed by CI metagenomics methods in the water samples examined (stream and reservoir) in the present study using 70 to 150 million reads. For these waters, CI data was insufficient for the monitoring aims of the NARMS program, but QMGS data met the challenge of reporting clinically important antimicrobial resistance from both water sources.

Important studies have described AMR in water and effluents [3-5, 26-36], but lack of a consensus set of tools to harmonize methodologies and analyses across multiple groups’ efforts (from water collection and laboratory processing to bioinformatic analysis and reporting) leaves us without a ready, common source of information to support local, national and global AMR monitoring of the environment. Metagenomic approaches provide valuable surveillance of species such as *Mycobacterium, Mycoplasma* and *Neisseria* (as shown in Figure 11), but for surveillance of NARMS critically important gene targets, metagenomics should be coupled with microbial pre-enrichment using relevant bacterial growth media.

Quasimetagenomic approaches have an established history of expediting whole genome sequencing (WGS) based source tracking and plasmid identification, and for describing multi-serovar diversity associated with outbreaks that might otherwise be overlooked [11, 12, 37]. Here we demonstrate that QMGS data also provides robust utility for AMR reporting from surface waters. With additional modifications to achieve greater microbiological selectivity, it should be possible to develop more expansive coverage of specific targets and consortia of targets. Even in enriched samples for Sligo and Patuxent, with an average of 70 million reads per replicate, coverage of taxa responsible for the most AMR deaths worldwide (*Escherichia coli*, *Staphylococcus aureus*, *Klebsiella pneumoniae*, *Streptococcus pneumoniae*, *Acinetobacter baumannii*, and *Pseudomonas aeruginosa* [22]) together comprised less than 6 % of total enriched data. This result, in contrast to less than 1% coverage in unenriched data, is a vast improvement but still leaves opportunity for further optimization.

The QMGS data presented here represents water incubated for 24 H at 37° in two culture broths, BPW and TSB. Biases were already evident in comparisons of BPW to TSB – for example, *Pseudomonas* species being predominantly observed in BPW and not in TSB enrichments. Another focus of our methods development was the comparison of 10 L and 50L of water to see if significant differences in coverage may be associated with either volume, but there were not clear differences suggesting the 50 L provides a significant increase in detecting capacity. There are many opportunities for advanced, precision QMGS targeting of important species and consortia of species by adjusting nutrients, oxygen tension, time, temperature, and addition of antibiotics. Bioinformatic methods also require optimization, validation, and standardization for harmonized reporting of taxonomy and resistance determinants. All bioinformatic approaches provided slightly to vastly different annotation and detection sensitivity for AMR associated with Sligo and Patuxent water.

The Centers for Disease Control and Prevention (CDC) estimate that millions of people in the United States are impacted by waterborne diseases every year [38]. The WHO reports that 144 million people rely on untreated surface water [39], with projections that by 2025, more than half of the world’s population will live in water stressed areas [39]. While the precise intersection of water and AMR morbidity and mortality is not fully understood, the fundamental importance of water to the cells of all living organisms is quite clear. Surface waters have the potential to serve as sentinels for global One Health monitoring of antimicrobial resistance and a wide range of anthropogenic pollutants that may correlate with risk factors. Murry et al. estimate there were 1.27 million deaths attributable to antimicrobial resistance in 2019 [22]. That number is close to the global number of HIV (680,000) and malaria (627,000) deaths combined [40–42]. Understanding the flow of antimicrobial resistance through aquatic ecosystems is a fundamental objective for One Health research and a new research priority for the National Antimicrobial Resistance Monitoring System.

As laboratory and informatic methods are harmonized in collaborative endeavors spanning microbiology, molecular biology, chemistry, hydrology, epidemiology, and interoperable metadata ontology, it will be possible to develop global resources to describe phenotype evolution, plasmid dynamics and an overall understanding of the flow of AMR through water and index it to resistance present in human and animal pathogens. Our findings suggest that metagenomic and quasimetagenomic data used together, provide a framework to support a new era of One Health monitoring and identification of emerging resistance.

## Materials and Methods

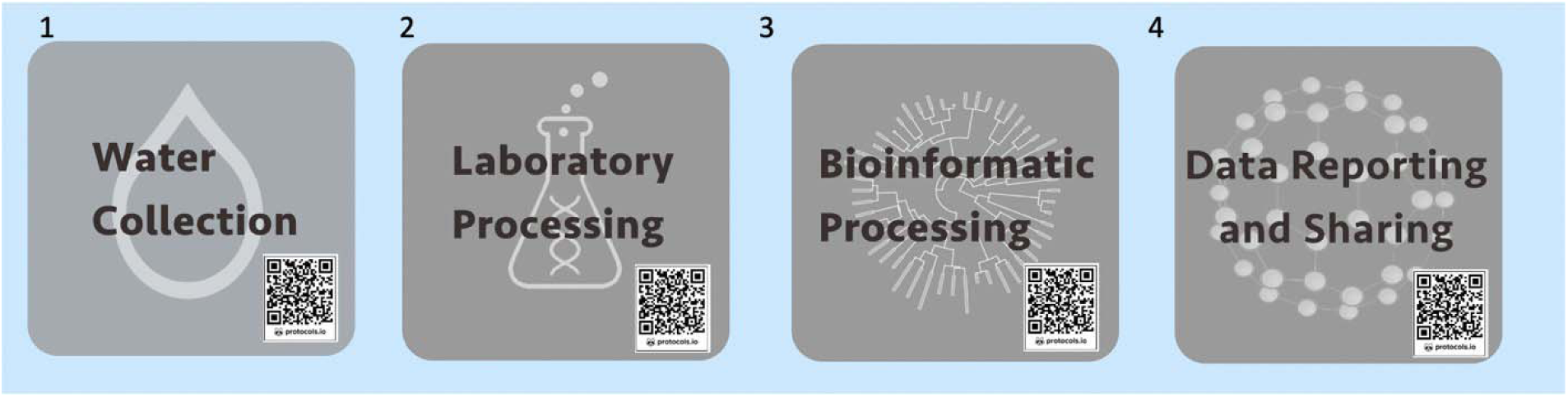

Each set of protocols used here is available with barcode scanning which links to open access visualized methodologies at: protocols.io covering every detail of water collection, laboratory processing, bioinformatic analysis and data reporting and sharing.

### Water collection

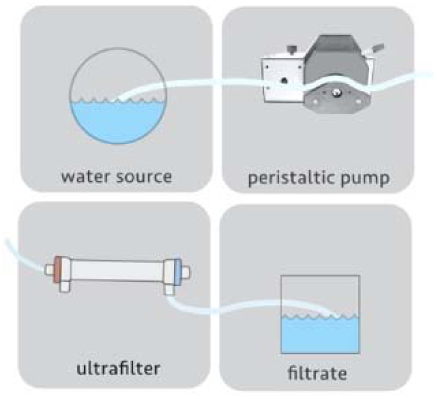

https://protocols.io/view/dead-end-ultrafiltration-water-collection-bztfp6jn

### Water Collection: Volume

Numerous studies have documented that increased water volumes correlate with increased likelihood of detecting bacterial pathogens but less work has described this correlation for the resistome. It stands to reason that important AMR phenotypes of clinically significant human and animal pathogens will be more likely to be recovered when more water is examined.

### Water Collection: DEUF

Dead end ultrafiltration (DEUF) at 10 and 50 L volumes was used to collect water from Sligo and Patuxent water sources[27]. DEUF was done using a Hemodialyzer Rexeed 25S filters (AsahiKasei, Chiyoda, Tokyo, Japan) and a Geopump peristaltic pump (Geotech, Denver, CO). Cells and particles are caught in the hollow fiber membranes within the ultrafilter filter, while filtrate passes through. Ultrafilter membrane separation collects particles between nano and micro (pore size range of 0.001–0.05 μm) and all manner of larger organisms and debris. The size range captured by ultrafiltration is ideal for examination of viruses, bacteria, fungi, and protists in water. After collection, filters are capped, bagged and stored at 4°C, until backflushing

### Laboratory Processing

#### Backflushing

First step for laboratory processing is the ‘backflushing’ of the filter.

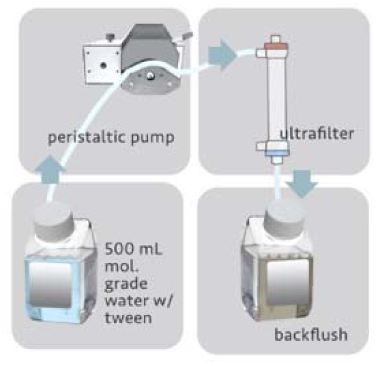

https://www.protocols.io/edit/backflush-of-dead-end-ultrafilter-bzuxp6xn

Backflush concentrate was used for culture independent (CI) metagenomics by direct DNA extraction of water filters passed through 0.2 micron filters. Nucleic acid extraction was conducted directly on the filters as part of the Qiagen DNeasy PowerWater DNA extraction protocol (Qiagen, Germantown, MD, United States) according to the manufacturer’s specifications (Qiagen PowerWater Kit Handbook).

#### Enrichment

Enrichment for quasimetagenomes was achieved by adding BPW and TSB at a 1 to 1 ratio (25 ml to 25 ml) (4 replicates of backflushed filtrate from each of 10L and 50L ultrafilter collections with incubation at 37 ° for 24 H). After 24 H of incubation at 37°, 2 ml aliquots were removed, centrifuged and DNA was extracted from the pellet using the Zymo High Molecular Weight DNA Extraction kit according to the manufacturers specifications (Zymo Quick-DNA Magbead Handbook).

### Library Preparation and Sequencing

DNA libraries from both CI and QMGS samples was prepared using the Illumina DNA Library Prep according to the manufacturers specifications (Illumina).

https://www.protocols.io/edit/illumina-dna-prep-sop-bzstp6en

Sequencing was performed on a NextSeq 500 according to the manufacturer’s specifications. All sequencing runs were performed in paired end mode with 2 x 150 cycles using the NextSeq 500/550 v2.5 High Output Kit (150 Cycles). Libraries were diluted to 1.8 pM according to the manufacturer’s specifications (NextSeq Denature and Dilute Libraries Guide).

### Bioinformatic Analyses

Files were demultiplexed (bcl to fastq) and screened/trimmed using Trimmomatic[43].

Four replicates of each treatment with reads per sample spanning 20 million to 150 million reads were used for further downstream analyses. Fastqs were run on the AMR++ pipeline[7] with the Megares database v2 using the CFSAN High Performance Cluster (HPC) with default parameters. All fluoroquinolones requiring SNP confirmation were verified by identifying the SNPs conferring resistance in Tablet [44] using the AMR++ output. For annotation by AMR FinderPlus, fastq files were aligned against the AMRFinder Plus database using SAUTE [45] on the CFSAN HPC. https://github.com/ncbi/amr/wiki/Methods. RGI and Blast were used with CARD[17] database to annotate sequence data according to default parameters. Reads were also evaluated using the COSMOS ID analytical pipeline (AMR database update July 2021 https://www.cosmosid.com). Counts and abundances from AMR annotation outputs were ‘normalized’ using scripts to assess ‘reads per kilobase of transcript’ (RPKM) to normalize gene reporting between different sites by accommodating for variation in number of sequencing reads per sample and gene length. Total reads in the sample were divided by 1,000,000 “per million” scaling factor to normalize for sequencing depth and provide ‘reads per million’ (RPM). RPM values were then divided by length of each gene in kilobases to report ‘RPKM. https://github.com/SethCommichaux/AMRplusplus

Files were demultiplexed (bcl to fastq) and screened/trimmed using Trimmomatic[43]. Four replicates of each treatment with reads per sample spanning 20 million to 150 million reads were used for further downstream analyses. Fastqs were run on the AMR++ pipeline[7] with the Megares database v2 using the CFSAN High Performance Cluster (HPC) with default parameters. All fluoroquinolones requiring SNP confirmation were verified by identifying the SNPs conferring resistance in Tablet [44] using the AMR++ output. For annotation by AMR FinderPlus, fastq files were aligned against the AMRFinder Plus database using SAUTE [45] on the CFSAN HPC.

#### Annotation of Metagenomic Outputs of Antimicrobial Resistance

Each pipeline used to examine Sligo and Patuxent data generated annotations by different algorithmic approaches and slightly different databases and output styles. AMRFinderPlus uses Hidden Markov Models (HMMs) with customized algorithms to identify AMR genes, point mutations, stress responses, and virulence genes. AMRFinderPlus can use both protein and nucleotide sequences. Outputs include gene length and contig position information for users to further their own evaluations of the annotations[14]. The pipeline was primarily designed for use with genomes. AMR++ was designed for use with large datasets of short read metagenomic data. It handles terabyte sized data fast and accurately for count-based data. The latest update in AMR++’s associated database, MEGARes 2.0 incorporates published resistance sequences (∼8,000 hand curated) for antimicrobial drugs, metal, and biocide resistance determinants [16]. CosmosID uses kmers which are excellent for detection (sensitivity) but sometimes lacking in specificity. While Cosmos’s approach and database is proprietary, there is very clear overlap with AMRFinderPlus annotation which uses the NCBI databases. For all pipelines used, greatly enhanced capacity to describe resistomes was observed with data from quasimetagenomic approaches[15]. The Comprehensive Antibiotic Resistance Database (CARD) (https://card.mcmaster.ca) provides highly curated reference sequences of both nucleotides and proteins with a highly structured ontology to support analysis and prediction[17]. Plasmids were annotated using ‘Platon’ according to the default parameters [46] (https://github.com/oschwengers/platon).

### Taxonomy

All taxonomic annotation was accomplished using an in-house FDA kmer pipeline developed at CFSAN, FDA, hand curated since 2014 to address FDA pathovars, using a high-performance computer cluster and also available on GalaxyTRAKR http://galaxytrakr.org.

## Supporting information

S Fig 1. Multibiocide resistance in Sligo and Patuxent water by CI and QMGS approaches

## Data Reporting and Sharing

### Reporting

Pipeline annotation outputs were visualized using R and Graphlan [47]. Replicates of water samples were merged for certain visualizations to simplify reporting.

### Data Sharing

All sequences have been deposited in NARMS Water: NARMS Water Metagenomes in the BioProject: SubmissionID: SUB10902319 BioProject ID: PRJNA794347 http://www.ncbi.nlm.nih.gov/bioproject/794347

## Definitions

**\*MTRAD** genes confer resistance to multi-drug resistance and include: ***asmA*** gene is hypothesized to control expression of the ***marRAB*** operon in *Salmonella enterica*, the ***nalD*** gene encodes a transcriptional regulator of the MexAB-OprM efflux system in *Pseudomonas aeruginosa*, the ***ramAR*** genes encode ***araC*** family transcriptional regulators that affect expression of the ***acrAB*** multi-drug efflux system, and ***ramAR*** genes encode ***araC*** family transcriptional regulators that affect expression of the ***acrAB*** multi-drug efflux system[16].

**ESKAPE8** The ESKAPE8 list used here is comprised of the 6 ESKAPE pathogens; *Enterococcus faecium, Staphylococcus aureus, Klebsiella pneumoniae, Acinetobacter baumannii, Pseudomonas aeruginosa, and Enterobacter spp* merged with only two not represented in that list from the top 6 species responsible for global AMR associated mortality ie: *Escherichia coli*, *Staphylococcus aureus*, *Klebsiella pneumoniae*, *Streptococcus pneumoniae*, *Acinetobacter baumannii*, and *Pseudomonas aeruginosa.* Thus ESKAPE8 = *Enterococcus faecium, Staphylococcus aureus, Klebsiella pneumoniae, Acinetobacter baumannii, Pseudomonas aeruginosa, Enterobacter spp., Escherichia coli* and *Streptococcus pneumoniae*.

## Acknowledgments

We would like to acknowledge the architects and scientists that support the CFSAN High Performance Cluster.

S Fig 1. Multibiocide resistance in Sligo and Patuxent water by CI and QMGS approaches.

