## Supplementary figures and images for "Advancing antimicrobial resistance monitoring in surface waters with metagenomic and quasimetagenomic methods"

### S Fig 1. Multibiocide resistance in Sligo and Patuxent water by CI and QMGS approaches

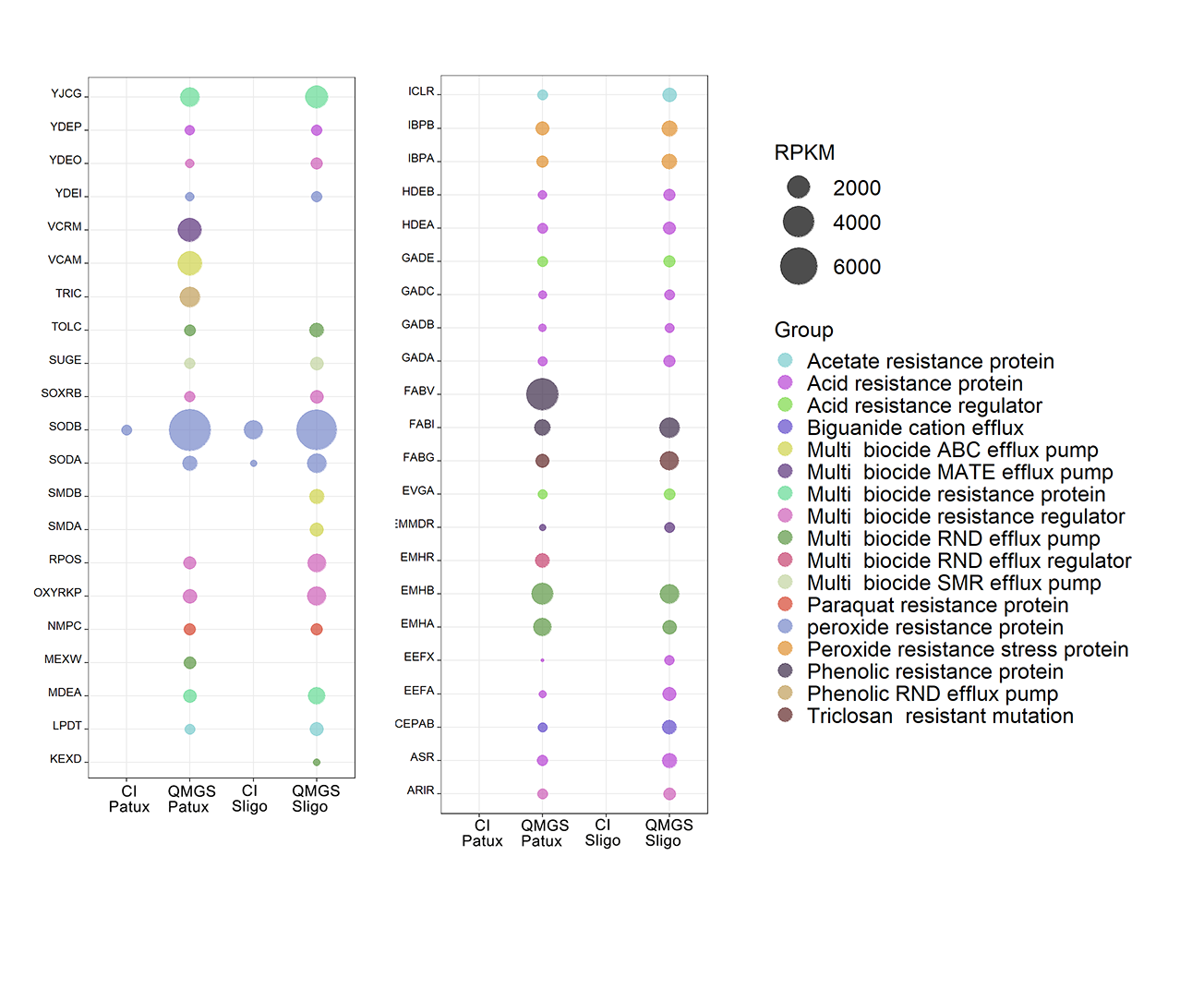
